# Impact of Microscopic Quantum Mechanisms on Macroscopic Epigenetic Regulation through Histone Deacetylation

**DOI:** 10.1101/2023.07.18.549607

**Authors:** Takeshi Yasuda, Tomoo Ogi, Nakako Nakajima, Tomoko Yanaka, Izumi Tanaka, Katsushi Tajima

## Abstract

The question of whether physical phenomena at a quantum level significantly impact aspects of macroscopic life has long remained unanswered. Histone modification by acetylation regulates the transcriptional activity of genes, and thereby broadly impacts cellular metabolism. In chemical reactions, the quantum tunneling effect is a phenomenon in which a small quantum particle of the reactant can pass through the potential energy barrier, even if it does not have sufficient energy to overcome the barrier. Here, we demonstrated that quantum effects are involved in the enzymatic reaction of histone deacetylation, by monitoring kinetic isotope effects due to hydrogen isotopes of water molecules and their temperature dependence as indicators. Due to the kinetic isotope effects associated with the quantum effects, the reaction rate balance between histone acetylation and deacetylation in cells was altered with heavy water, which changed epigenetic transcription regulation in the cells. Thus, microscopic quantum mechanisms exist in histone deacetylation, thereby broadly impacting macroscopic life phenomena through epigenetic regulation.

**Teaser:** Quantum effects in enzymatic reaction of histone deacetylation latently influence life phenomena through epigenetic regulation.

## Introduction

In eukaryotes, including humans, genomic DNA is wrapped around histone proteins to form nucleosomes, which are further folded into chromatin, and thereby the DNA is accommodated within the cell (*1*). Histone proteins form an octamer containing two each of the four protein subunits, H2A, H2B, H3, and H4. Chromatin structures inhibit the access and movement on DNA of DNA binding proteins, and thereby impact diverse DNA-mediated transactions. The inhibitory effects of chromatin structure and their release by its conformational changes regulate DNA metabolism, including gene transcription (*2*). The amino-terminal region of each histone subunit is a positively charged lysine-rich region called the histone tail, which is involved in binding to negatively charged DNA. Lysines in the histone tail undergo various protein modifications, including acetylation. The loss of the positive charge of lysine by acetylation weakens histone binding to DNA and converts the condensed chromatin to a relaxed, open chromatin structure. The conversion of chromatin structure via histone acetylation activates transcription by promoting the recruitment of transcription enzymes to genes and their movement on genes (*1*-*5*). Histone acetylation modifications are catalyzed by histone acetylases (HATs), while histone deacetylases (HDACs) remove these modifications (*6*). The regulation of histone acetylation levels via HATs and HDACs is important for the precise control of gene expression.

The regulation mechanisms of gene expression via chromatin structure and histone modifications have been elucidated from the viewpoints of genetics, biochemistry, and cell biology. In contrast, quantum biology aims to elucidate life phenomena from a quantum theoretical perspective (*7*). Photons and electrons, which possess both particle and wave properties, are quanta. Therefore, the reaction mechanisms of enzymes that perform photosynthesis or DNA photorepair in response to photons have been studied from a quantum mechanical perspective (*8*-*10*). Quantum mechanics is involved in the cleavage and formation of chemical bonds mediated by electrons. Accordingly, quantum mechanics is also expected to function in reactions by enzymes that catalyze such chemical reactions. For a chemical reaction to proceed, the reactants in the ground state must become excited and exceed the potential energy required for the reaction (*11,12*). However, in quantum mechanics, if the temperature of the reactant is low and the reactant cannot exceed the potential energy, quantum particles of the reactant can pass through the potential energy barrier without reaching the transition state. This phenomenon is called quantum tunneling. The probability of this occurring decreases as the mass of the reactant increases. Quantum tunneling occurs in reactions involving electrons and hydrogen, with small atomic weights, but not atoms with large atomic weights (*11, 12*). Therefore, the probability of quantum tunneling is higher for hydrogen (H), with an atomic weight of 1, than for deuterium (D), an isotope of hydrogen with an atomic weight of 2, resulting in a faster chemical reaction rate for H than for D. The change in reaction rate that occurs when one of the atoms of a reactant is substituted by an isotope is called the kinetic isotope effect. Thus, the kinetic isotope effect due to H and D is related to the quantum tunneling effect. When the reaction temperature is high and the reactant can surmount the potential energy barrier without quantum tunneling, the kinetic isotope effect is reduced or eliminated (*11*-*14*). By using the kinetic isotope effect and its temperature dependence as an indicator for quantum tunneling, its influences have been demonstrated for the enzymatic chemical reactions of alcohol dehydrogenases and similar family proteins that catalyze hydrogen-transfer reactions during substrate oxidation (*15, 16*). However, little is known about the involvement of quantum tunneling effects in proteins participating in various other life phenomena and their effects on cells. In this study, we investigated whether the quantum tunneling effect is involved in another type of important enzymatic reaction, the deacetylation of histone proteins. In addition, we assessed the influence of the quantum tunneling effect on gene expression in cells.

## Results

### Concept of this research method

Chemical reactions of quantum particle reactants at the ground state (GS) can proceed without reaching the transition state (TS) due to the quantum tunneling effect (fig. S1A). As described in textbooks such as Atkins’ Physical Chemistry, the probability of the quantum tunneling effects of a quantum particle is derived from the Schrödinger equation, the fundamental equation in quantum mechanics, in the one-dimensional box barrier model (fig. S1B), and is shown by the following formulas (*11*).

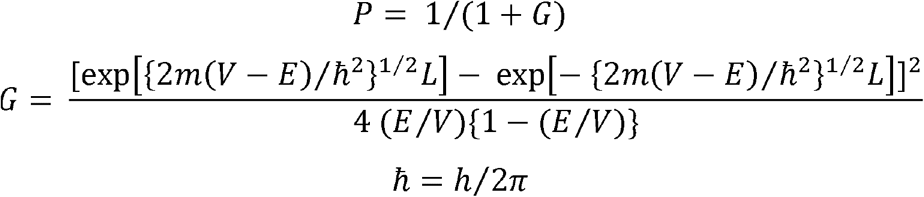

Where *P* is the probability of the quantum tunneling effect, *V* is the potential energy of the particle, *E* is its total energy, *m* is the mass of the particle, and *h* is Planck’s constant. Since the quantum tunneling effect is significant where *E* < *V*; for example, when *E*/*V* is assumed as 0.5, then 4(*E*/*V*){1-(*E*/*V*)} = 1, and 2*m*(*V*-*E*) = *m*. The formula for *G* thus simplifies to

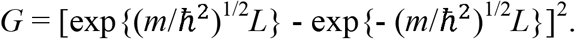

Comparing the inside of this formula between ^1^H and ^2^H,

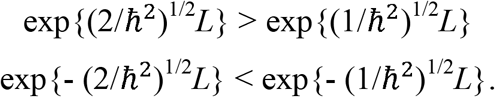

Thus,

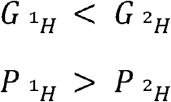

These formulas clarify that hydrogen has a higher probability of quantum tunneling than deuterium, and that the kinetic isotope effect occurs via the different probabilities of quantum tunneling between hydrogen and deuterium. In addition, as described in Atkins’ Physical Chemistry, the probability of quantum tunneling (*P*) decreases exponentially with *m*^1/2^. Therefore, quantum tunneling is more difficult for heavier particles (*12*). At higher reaction temperatures, E > V, the differences in quantum tunneling effects due to differences in particle mass disappear. Therefore, the kinetic isotope effect due to quantum tunneling is temperature dependent.

As described in Atkins’ Physical Chemistry (*12*), the kinetic isotope effect is also caused by differences in the zero-point vibrational potential energies of the chemical bonds involved in the reaction, which are likewise related to quantum mechanics (fig. S1C). For example, the ratio of the reaction rate constants between the cleavage of O-H and O-D bonds, *k*(O-D)/*k*(O-H), is shown by the following formulas (*12*).

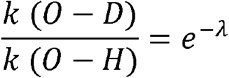

with

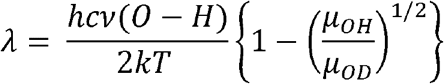

Where *h* is Planck’s constant, *c* is the speed of light, ν(O-H) is the vibrational wavenumber of the O-H bond, μOH and μOD are the relevant effective masses, *k* is the Boltzmann constant, and *T* is the absolute temperature. Therefore, the kinetic isotope effect is caused by the differences in the relevant effective masses between OH and OD. In addition, the formulas indicate that *k*(O-D)/*k*(O-H) depends on temperature. However, comparing k(O-D)/k(O-H) at 15°C (288K) and 42°C (315K), e^-λ^ (at 315 K)/e^-λ^ (at 288 K) = 1.000297663340558. Thus, in the temperature range used in our experiments, the isotope effects caused by the differences in zero-point vibrational potential energy are less sensitive to temperature variation.

The enzymatic chemical reaction of deacetylation, which removes acetyl groups from acetylated histones, is a hydrolytic reaction that directly involves water molecules in the reaction solution (*17*-*19*). A hydrogen atom in the water molecule attacks the chemical bond of the acetylated lysine to remove the acetyl group, and binds to the site from which the acetyl group has been removed (fig. S1D). When heavy water (D_2_O) is used instead of water (H_2_O) in the reaction solution, the deuterium (D) of D_2_O binds to the lysine. Therefore, by comparing the reaction rates of histone deacetylation in the presence of H_2_O or D_2_O and their temperature dependence *in vitro*, it is possible to investigate whether quantum tunneling effects are involved in the enzymatic chemical reaction of histone deacetylation.

### Kinetic isotope effects on SIRT3-mediated histone deacetylation

To examine the kinetic isotope effects on histone deacetylation, we used p300, a human HAT, SIRT3, a human HDAC, and unmodified recombinant histone octamer proteins as their substrates (*20, 21*). At first, histone octamer proteins are incubated with p300 and ^14^C-labeled Acetyl Coenzyme A (Ac-CoA) in reaction buffer containing H_2_O or D_2_O, for histone acetylation at 30°C (HAT assay). A portion of each reaction mixture containing the acetylated histones is then incubated with SIRT3 and NAD^+^ in reaction buffer containing H_2_O or D_2_O, respectively (HDAC assay) (Fig. 1A). Each sample before and after the HDAC assay is separated by sodium dodecyl sulfate (SDS)-polyacrylamide gel electrophoresis (PAGE), and detected with Coomassie Brilliant Blue (CBB) staining and autoradiography. The total proteins are stained by CBB, and the acetylated proteins are detected by autoradiography. The relative quantities of the mixture of histones H3, H2A, and H2B (top bands), and H4 (bottom bands) are quantified. The kinetic isotope effects were detected at 15°C (Fig. 1 B-E), but disappeared at 42°C (Fig. 2). Therefore, the temperature dependency of the kinetic isotope effects suggests the involvement of the quantum tunneling effect in the SIRT3-mediated deacetylation of histone proteins.

**Fig. 1.**
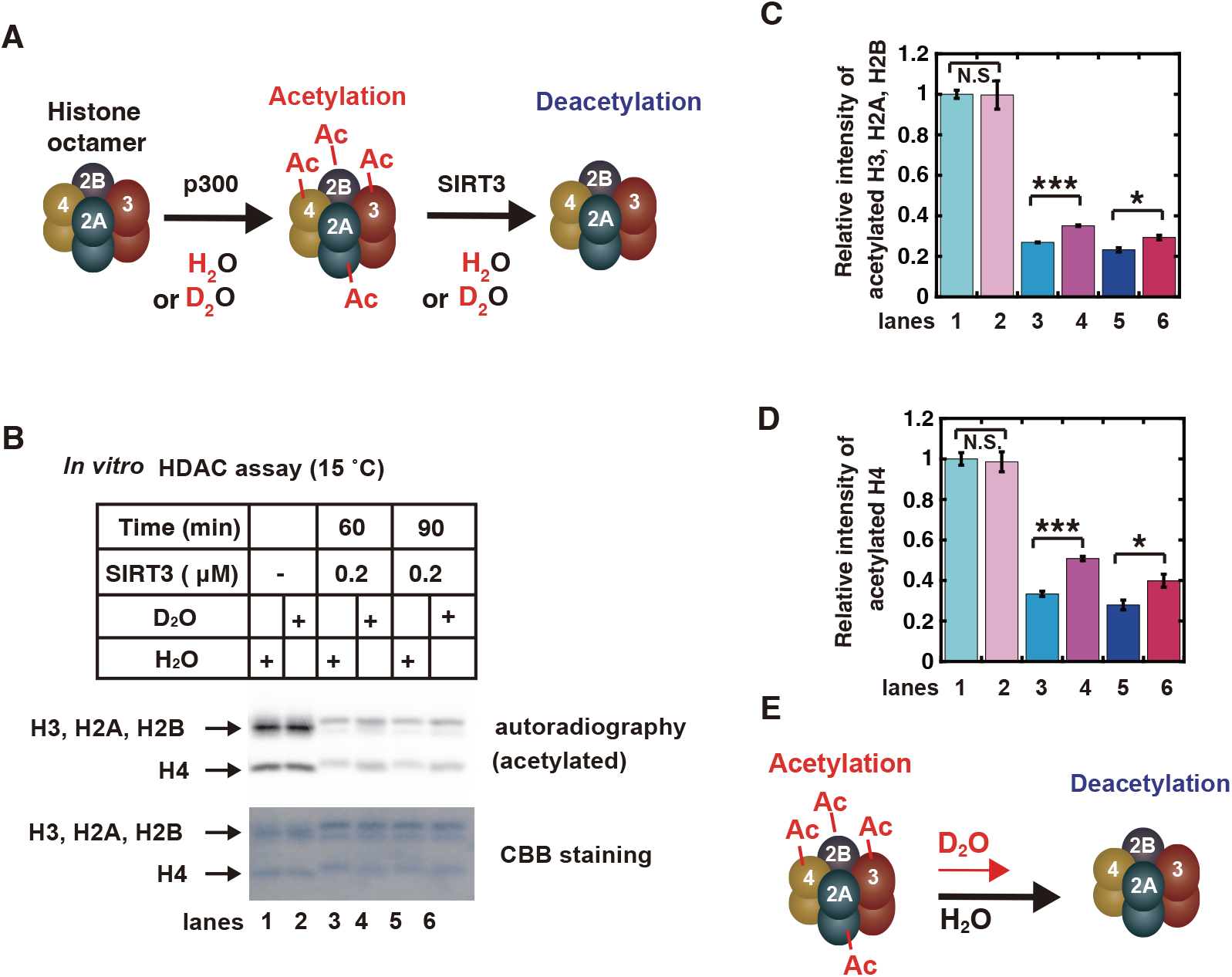
Kinetic isotope effect on histone protein deacetylation reactions. (**A**) Schematic representation of the histone deacetylation assay. (**B**) *In vitro* acetylation and deacetylation assays of histone proteins were performed, as described in the Materials and Methods. The deacetylation reactions were performed with the indicated amounts of SIRT3 at 15°C for the indicated times. The reaction mixtures were subjected to SDS-PAGE, followed by CBB staining (bottom) and autoradiography (top, acetylated proteins). (**C** and **D**) The relative band intensities of acetylated H3, H2A, and H2B (C) and acetylated H4 (D). The graph shows the mean values and standard errors of the mean from 3 independent experiments. The samples connected by lines were compared (\**P* <0.05, \*\*\**P* <0.001 and N.S., not significant by an unpaired Student’s t-test). (**E**) Schematic representation of the experimental results about the kinetic isotope effect on protein deacetylation reaction.

**Fig. 2.**
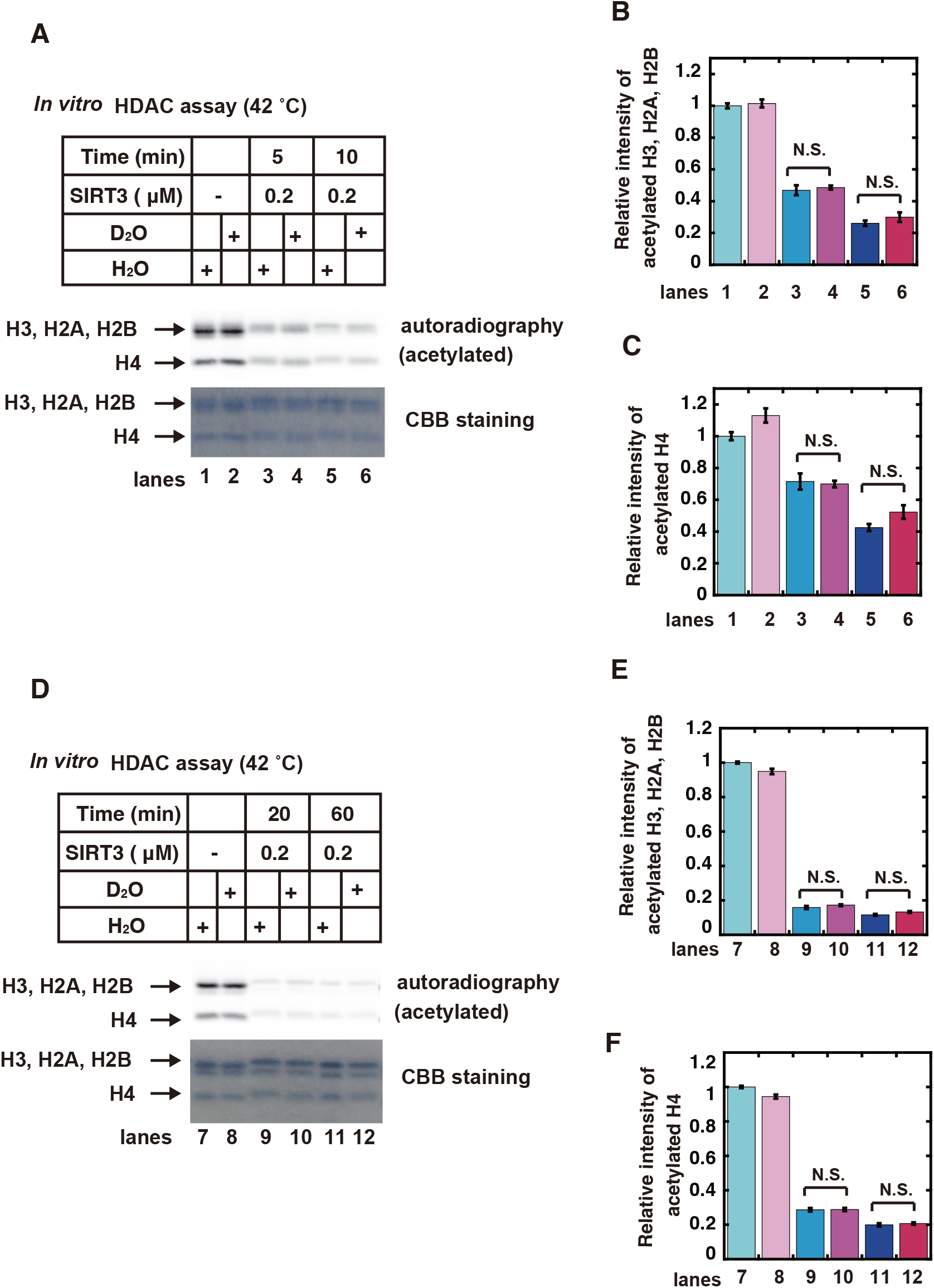
Temperature dependency of the kinetic isotope effect on SIRT3-mediated histone deacetylation. (**A** and **D**) *In vitro* acetylation and deacetylation assays of histone proteins were performed, as described in the Materials and Methods. The deacetylation reactions were performed with the indicated amount of SIRT3 at 42°C for the indicated times. The reaction mixtures were subjected to SDS-PAGE, followed by CBB staining (bottom) and autoradiography (top, acetylated proteins). (**B, C, E**, and **F**) The relative band intensities of acetylated H3, H2A, and H2B (B and E) and acetylated H4 (C and F) are shown in the graph. Mean values and standard errors of the mean from 3 independent experiments were plotted. The samples connected by lines were compared (N.S., not significant by an unpaired Student’s t-test).

### Isotope effect by D_2_O on histone acetylation and transcription in cells

Histone acetylation in cells is altered by the balance between acetylase and deacetylase activities, which are involved in transcriptional regulation (Fig. 3A) (*1*-*4*). Therefore, when histone deacetylases are inhibited, histone acetylation is induced, and thereby transcription is increased (Fig. 3A). Consistent with the kinetic isotope effect on the *in vitro* histone deacetylation, the induction of histone acetylation was detected when cells were exposed to D_2_O (Fig. 3B). A global transcriptional analysis with RNA-seq revealed that overall transcription changed much more by a D_2_O-treatment than by an siRNA-treatment against SIRT3 (Fig. 3C, fig. S2, and fig. S3). On average, the D_2_O-treatment increased transcription far more than the siRNA-treatment against SIRT3 in cells. The RNA-seq data analysis revealed that there were similarities in the groups of genes with transcription altered by D_2_O-treatment under both the mock and siRNA-treatment conditions (fig. S4, fig. S5, and fig. S6). The gene expressions from the RNA-seq data were analyzed (*22, 23*) and are shown in color on KEGG pathway maps (*24*) (figs. S7 to S10). The D_2_O treatment induced the expression of genes involved in cellular defenses and cytokines (fig. S7 A and B, and fig. S10B), but decreased the transcription of genes involved in DNA transactions and the cell cycle (fig. S8 A and B, and fig. S9 A and B). Among the genes involved in homologous recombination (HR), the expression of most of the genes required for HR repair, such as MRN complexes (*25, 26*), was decreased by D_2_O-treatment, whereas the expression of SYCP3 (*27*), which suppresses HR repair, was increased (fig. S9B). The expression of genes regulated by p53, such as p21 and GADD45 (*28*), which suppress cell cycle progression, and PUMA and Noxa (*28*), which induce apoptosis, a mechanism of programmed cell death, was increased by D_2_O-treatment (fig. S9A, and fig. S10 A and B). Conversely, the expression of Bcl2 (*28*), which suppresses apoptosis, was decreased by D_2_O-treatment (fig. S10A). These effects of D_2_O on cellular transcription were similar to those of chemical HDAC inhibitors, which induce the expression of p21, GADD45, and pro-apoptotic genes and reduce the expression of pro-survival and cell cycle genes (*6, 29*).

**Fig. 3.**
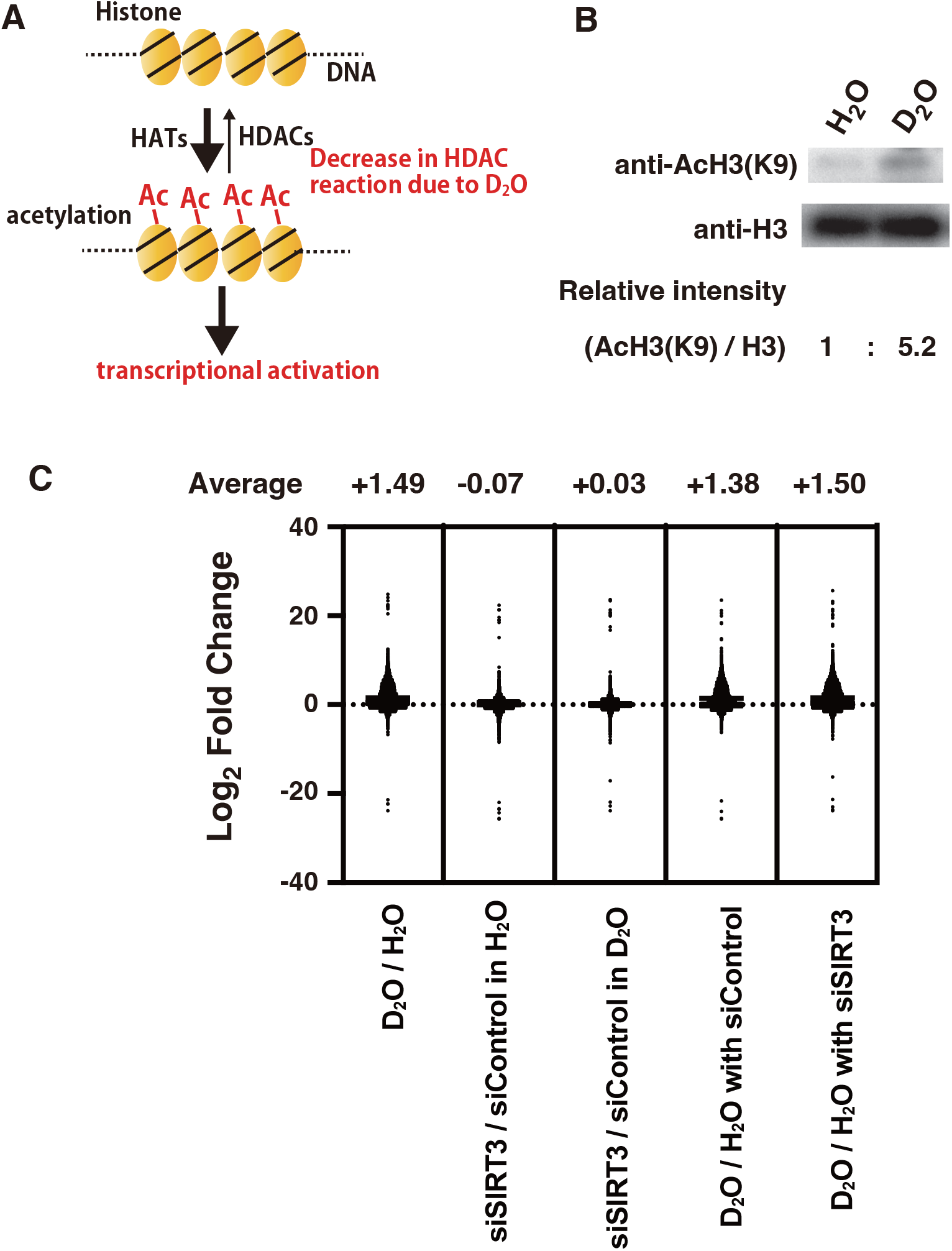
Isotope effects of D_2_O on histone acetylation and transcription in human cells. (**A**) Schematic representation of the regulation of histone acetylation-mediated transcriptional activation by HATs and HDACs and the isotope effect by D_2_O. (**B**) U937 human cells were cultured in medium containing H_2_O or D_2_O for 2 h. The cell extracts were subjected to immunoblotting analyses with the indicated antibodies. The relative band intensities of acetylated H3 at K9 (AcH3(K9)) bands normalized to those of the H3 bands are shown below the immunoblots. (**C**) RNA-seq analyses were performed with HeLa pDR-GFP human cells cultured in medium containing H_2_O or D_2_O for 5 h, as described in the Materials and Methods. The dots in the graph show the mean values of Log_2_ fold change of each gene from 6 samples. The average values of Log_2_ fold change of whole genes are shown in each of the comparison conditions.

### Isotope effect by D_2_O on p53-dependent apoptosis and necrosis

The RNA-seq data revealed that D_2_O-treatment induced the expression of genes related to apoptosis induction in human cells. Consistently, apoptosis was actually induced by D_2_O-treatment in human cells (Fig. 4A). Another cell death pathway, necrosis, was also activated by D_2_O-treatment (Fig. 4B). In addition, since both apoptosis and necrosis were decreased by about half in a p53-null background (Fig. 4 C and D), both cell death pathways induced by D_2_O were partially dependent on p53. This p53 dependency is consistent with the RNA-seq analysis results.

**Fig. 4.**
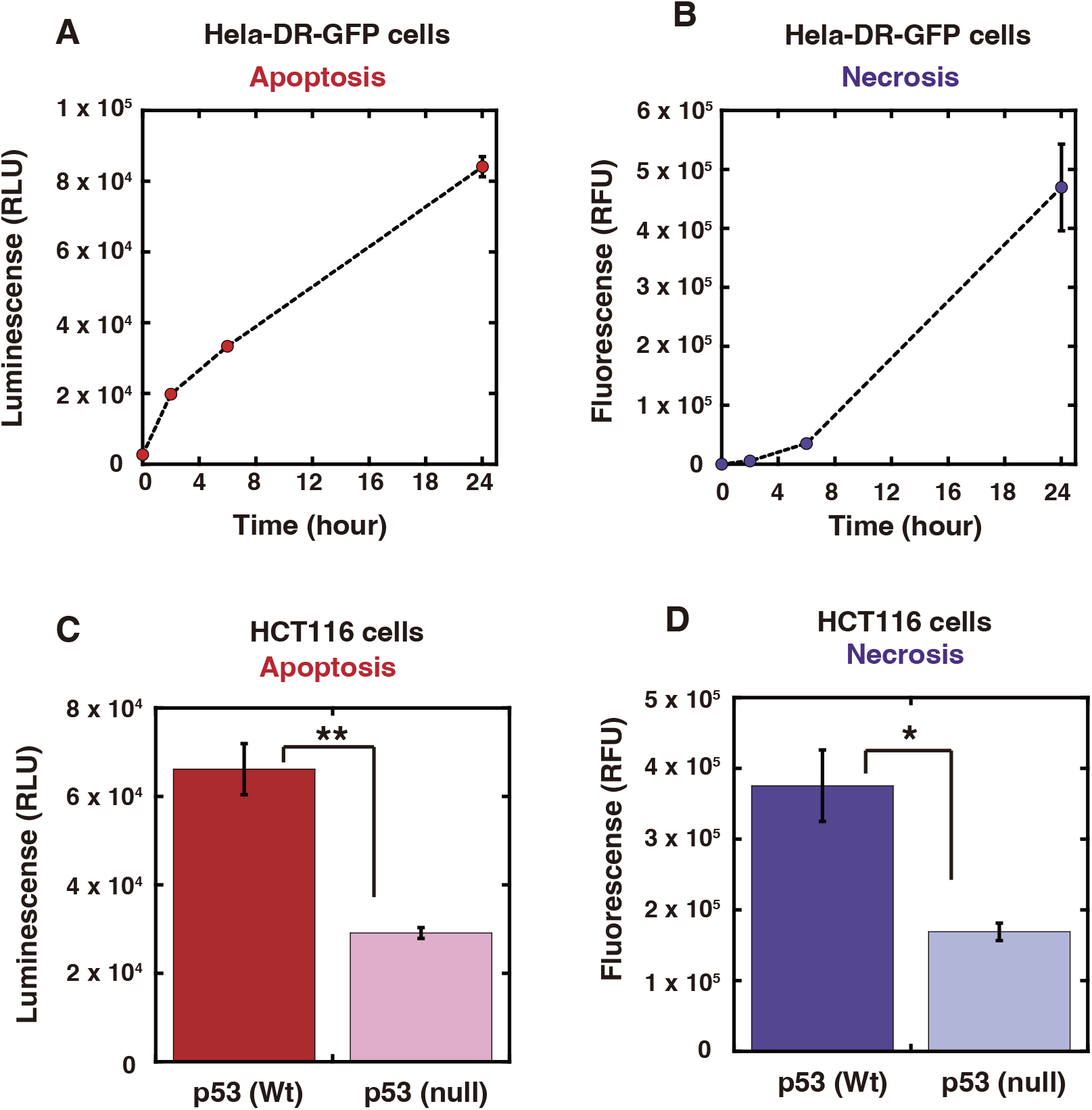
Apoptosis and necrosis inductions by D_2_O. (**A** and **B**) HeLa pDR-GFP cells were cultured in medium containing 100% D_2_O for the indicated times. (**C** and **D**) HCT116 (p53 wild type or null) cells were cultured in medium containing 100% D_2_O for 24 h. (**A** to **D**) Apoptosis and necrosis assays were performed, as described in the Materials and Methods. The graphs show the mean values and standard errors of the mean from 3 (A and B) or 4 (C and D) samples. (**C** and **D**) The samples connected by lines were compared (\**P* <0.05 and \*\**P* <0.01 by an unpaired Student’s t-test).

## Discussion

Quantum biology is expected to solve the mysteries of life phenomena that cannot be deciphered by traditional life sciences (*7, 30*). Several interesting examples of quantum biology have been introduced, such as the quantum magnetic compass of migratory birds and the quantum mechanism of efficient photosynthesis (*7, 30*). However, since these quantum biological research topics are still limited in scope, the significance of the quantum-level effects in life phenomena has remained elusive. In this study, using kinetic isotope effects and their temperature dependence as indicators, we show for the first time the involvement of quantum effects in the enzymatic chemical reaction catalyzing histone protein deacetylation. Since various DNA-mediated transactions are regulated by the level of histone acetylation, quantum effects on histone deacetylation reactions are expected to have a significant impact on a wide variety of life phenomena.

Histone deacetylation by SIRT3 exhibited a kinetic isotope effect, in which the reaction rate was slower with D_2_O than with H_2_O in the reaction solution. In relation to this *in vitro* quantum effect, an *in vivo* isotope effect was observed in which the exposure of human cells to D_2_O induced cellular histone acetylation. The addition of histone deacetylase inhibitors to cells induces histone acetylation because the intracellular acetylation equilibrium shifts in the direction of acetylation induction by HATs. The effect of D_2_O on intracellular histone acetylation is also presumably due to a change in the histone acetylation balance, by decreasing the deacetylation reaction rate. In relation to the fact that the induction of histone acetylation activates transcription, the addition of D_2_O resulted in an overall activation of intracellular transcription. In contrast to these effects by hydrogen isotopes, carbon and nitrogen isotopes, which have large atomic weights and thereby their quantum tunneling effects are not important, have almost no effect on the cells (*31*). Considering these facts, the cellular effects by D_2_O should be the isotope effects at the quantum level. Eighteen deacetylases, including SIRT3, have been identified in human cells. Since the knockdown of SIRT3 alone had no remarkable effect on cellular transcription, in contrast to the effects of the D_2_O-treatment, we speculate that D_2_O may have a similar isotope effect on other deacetylases involved in hydrolysis, and thereby exert a significant impact on overall cellular transcription.

Similar to the induction of apoptosis in cancer cells by histone deacetylase inhibitors (*6, 29*), D_2_O-treatment also induced apoptosis. Although D_2_O reportedly induces cell death (*32*-*34*), until now the underlying cause has not been discussed experimentally in relation to quantum effects. From the results of this study, we speculate that one of the reasons for the D_2_O-induced apoptosis is the reduction of the histone deacetylation reaction rate due to quantum effects. The induction of cell death by D_2_O was partially mediated by p53, a factor involved in the DNA damage stress response. Besides the p53-mediated cellular stress response, a variety of cellular stress responses involved in defense functions were also induced by D_2_O-treatment. Since the stress on cells mediated by the kinetic isotope effects of deuterium is thought to be related to quantum effects, we refer to this stress response as the quantum stress response.

Thus, the appearance of the isotopic effects of heavy water is due to the fact that the quantum effects are indeed present in hydrolytic enzymatic reactions. Presumably, in the time flow of life phenomena, enzymatic chemical reaction rates have been optimized for hydrogen of atomic weight 1, which constitutes ∼99.972% of the hydrogen atoms on earth (*35*). However, in an environment where deuterium is the only hydrogen present, the kinetic isotope effect of deuterium would reduce the reaction rate of enzymes, thus upsetting the balance of enzyme chemical reaction rates in the cell. This distortion of the reaction rate balance would cause significant stress, resulting in cell death.

## Materials and Methods

### Recombinant proteins and reagents

The recombinant human p300 protein was purified as described previously (*36*). Commercially purchased recombinant SIRT3 protein (#50014, Lot 2003, BPS Bioscience, San Diego, CA, USA) was used as described previously (*36*). Recombinant human histone H2A/H2B/H3.1/H4 octamer proteins were kindly provided by Dr. H. Kurumizaka (*21*). The following reagents were used: Acetyl Coenzyme A (A2181, Merck & Co., Inc., Whitehouse Station, NJ, USA), NAD^+^ (N7004, Merck & Co., Inc.), H_2_O (06442-95, Nacalai Tesque Inc., Kyoto, Japan), and D_2_O (D214H, Nacalai Tesque Inc.).

### Immunoblotting and antibodies

Immunoblotting analyses were performed as described previously (*36*). The following antibodies were used: anti-acetyl Histone H3 (Lys9) (06-942, Merck & Co., Inc.), and anti-GAPDH (2118, Cell Signaling).

### *In vitro* acetylation and deacetylation assays

*In vitro* acetylation assays of histone proteins were performed by incubating p300 (0.567 _μ_M) and human histone H2A/H2B/H3.1/H4 octamer proteins (4.62 _μ_M) in HAT buffer [50 mM Tris-HCl, 1 mM EDTA, 10% glycerol, 1 mM DTT, pH 8.0], made with H_2_O (06442-95, Nacalai Tesque Inc.) or D_2_O (D214H, Nacalai Tesque Inc.), in the presence of ^14^C-labeled Ac-CoA (170 _μ_M) at 30°C for 60 min. Subsequently, *in vitro* deacetylation assays of the acetylated histone proteins were performed by incubating an aliquot of the reaction mixture containing the acetylated histone octamer proteins (0.67 _μ_M) with the indicated amount of recombinant SIRT3 protein in HDAC buffer [25 mM Tris-HCl, 137 mM NaCl, 2.7 mM KCl, 1 mM MgCl_2_, 0.1 mg/ml BSA, pH 8.0], made with H_2_O (06442-95, Nacalai Tesque Inc.) or D_2_O (D214H, Nacalai Tesque Inc.), in the presence of 500 μM NAD^+^ at the indicated temperature. The reactions were fractionated by SDS-PAGE, and the gels were stained with Coomassie Brilliant Blue. The dried gels were photographed using an EOS 6D digital camera (Cannon, Tokyo, Japan) equipped with an EF 24-70 mm F4L IS USM lens (Cannon), and exposed to an imaging plate (Fuji Film, Tokyo, Japan). Radioisotopic images in the exposed imaging plate were analyzed by an FLA-3000 fluorescent image analyzer (Fuji Film).

### Cell culture

HeLa pDR-GFP cells, obtained from Dr. M. Jasin (*37*), were cultured in minimum essential medium (MEM) (1030700, Thermo Fisher Scientific, Waltham, MA, USA), supplemented with 10% fetal bovine serum (FBS), 2 mM L-glutamine, and 1% penicillin-streptomycin (PS). For experiments comparing the kinetic isotope effects due to H_2_O or D_2_O in these cells, the MEM solutions were prepared by dissolving MEM powder (61100, Thermo Fisher Scientific) in H_2_O (06442-95, Nacalai Tesque Inc.) or D_2_O (D214H, Nacalai Tesque Inc.) according to the manufacturer’s instructions.

U937 and HCT116 (p53 wild type or null) cells were cultured in RMPI 1640 medium (11875, Thermo Fisher Scientific) supplemented with 10% FBS and 1% PS. For experiments comparing the kinetic isotope effects due to H_2_O or D_2_O in the cells, the medium was prepared by dissolving RMPI 1640 powder (31800, Thermo Fisher Scientific) in H_2_O (06442-95, Nacalai Tesque Inc.) or D_2_O (D214H, Nacalai Tesque Inc.).

### siRNA treatments

Stealth Select siRNAs were purchased from Thermo Fisher Scientific. Stealth Select siRNA, HSS118726 (5’-AAUCAGCUCAGCUACAUCCUGCAGG-3’), was used for the siRNA treatment against SIRT3. Stealth RNAi negative control siRNA oligonucleotides (Thermo Fisher Scientific) were used for the negative controls. Lipofectamine RNAiMAX was used for the transfection of Stealth Select siRNAs into cells, according to the manufacturer’s transfection protocol.

### RNA extraction and RNA-seq analysis

Hela pDR-GFP cells (5×10^5^ cells/well in a 12-well plate) were untreated or transfected with siRNA. At 48 h after the siRNA-transfection, the cell culture medium was replaced with fresh medium. On the next day, the cell culture medium was changed to fresh culture medium made with H_2_O (06442-95, Nacalai Tesque Inc.) or D_2_O (D214H, Nacalai Tesque Inc.). After a 5 h incubation in the medium containing H_2_O or D_2_O, the cells were harvested by trypsinization and total RNA was isolated, using an RNeasy Mini Kit (Qiagen) according to the manufacturer’s instructions.

RNAseq was performed to investigate gene expression levels. mRNA was obtained from total RNA using the NEBNext Poly(A) mRNA Magnetic Isolation Module (NEB). The MGIEasy RNA Directional Library Prep Set (MGI Tech, Shinsen, China) was used to adjust the library. The libraries were sequenced on the DNBSEQ-G400 platform (MGI Tech), with paired-end flow cells to obtain 100–150 base pair reads with 100–200-fold coverage.

The obtained read counts data from RNA-seq experiments were analyzed with iDEP96 (*22*). The heatmap was generated using the following criteria with iDEP96: distance– correlation, linkage–average and cut-off Z score–4. The KEGG pathway maps (*24*) were colored based on the results of KEGG Pathview rendering (*23*) by iDEP96. The fold-change (log2) cutoff in the color code is 2. Permission has been obtained from Kanehisa laboratories for using KEGG pathway map images.

### Apoptosis and necrosis assays

For apoptosis and necrosis assays, cells were seeded in 96-well flat-bottom white plates (136101, Thermo Fisher Scientific) and 96-well flat-bottom black plates (137101, Thermo Fisher Scientific) at 10,000 cells/well. After 24 h, the cell culture medium was exchanged with fresh culture medium made with H_2_O (06442-95, Nacalai Tesque Inc.) or D_2_O (D214H, Nacalai Tesque Inc.). Apoptosis and necrosis assays were performed with a RealTime-Glo Annexin V Apoptosis and Necrosis Assay kit (JA1011, Promega, Madison, WI, USA) at the indicated times after changing the culture medium, using an ARVO X5 microplate reader (Perkin Elmer).

## Statistical analysis

The KaleidaGraph software, version 5.0, was used for statistical analyses. Statistical analyses for multiple comparisons were performed using a one-way ANOVA, as described previously (*38*). Statistical analyses between the data of two groups were performed using an unpaired Student’s t-test, as described previously (*38*).

## Acknowledgments

We thank Dr. M. Jasin (Memorial Sloan-Kettering Cancer Center, NYC, USA) for the Hela pDR-GFP cells. We deeply appreciate Dr. S. Hirayama (Emeritus Professor at Kyoto Institute of Technology, Kyoto, Japan) and Dr. H. Kitoh-Nishioka (Kindai University, Osaka, Japan) for helpful instructions and discussions about quantum tunnelling and kinetic isotope effects, and Dr. H. Kurumizaka and his laboratory members (The University of Tokyo, Tokyo, Japan) for purified histone proteins. This research was supported by Research Support Project for Life Science and Drug Discovery (Basis for Supporting Innovative Drug Discovery and Life Science Research (BINDS)) from AMED under Grant Number JP22ama121009 (H. Kurumizaka).

## Author contributions

Examples:

Conceptualization: T. Yasuda

Investigation: T. Yasuda, N.N., T.O., T. Yanaka, I.T.

Visualization: T. Yasuda, N.N., and T. Yanaka

Funding acquisition: T. Yasuda, K.T.

Writing—original draft: T. Yasuda

Writing—review & editing: T.O., K.T.

## Data and materials availability

All data are available in the main text or the supplementary materials. The biological materials are available upon request.

## Supplementary Materials

Figs. S1 to S10

Data S1 to S2

References

