## Supplemental Materials for "Impact of Microscopic Quantum Mechanisms on Macroscopic Epigenetic Regulation through Histone Deacetylation"

Science Advances
AAAS

### Supplementary Materials for

#### Impact of Microscopic Quantum Mechanisms on Macroscopic Epigenetic Regulation through Histone Deacetylation

Takeshi Yasuda *et al.*

**This PDF file includes:**

Figs. S1 to S10
Data S1 to S2
References 1 to 2

**Other Supplementary Materials for this manuscript include the following:**

Data S1 to S2

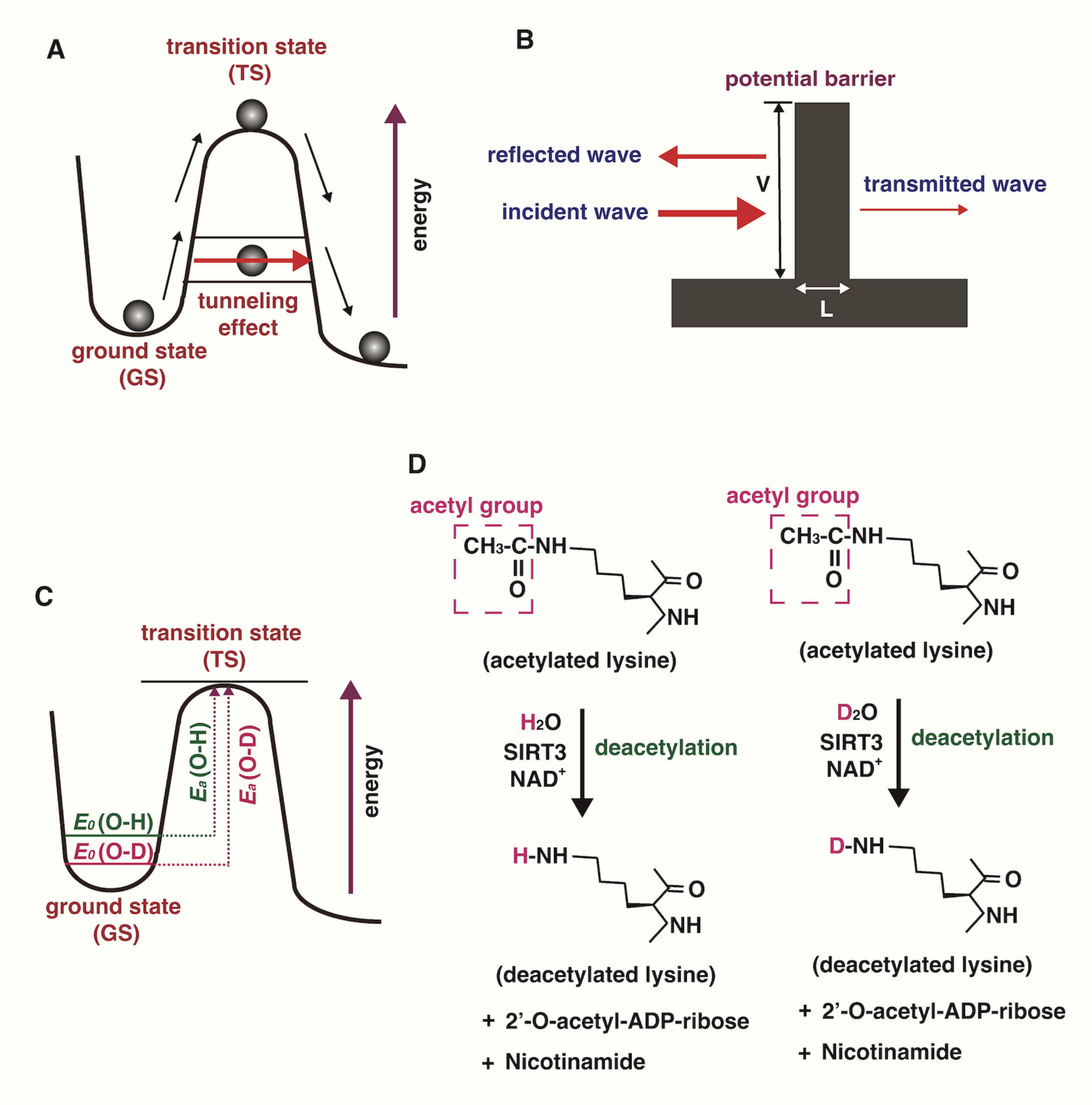


**Fig. S1.**

**Schematic representations of the concepts of this research method.** (**A**) Conceptual image of quantum tunneling. In quantum tunneling, the chemical reaction of the reactant (circle) at the ground state can proceed without reaching the transition state by penetrating through the potential energy barrier. (**B**) Conceptual diagram of quantum tunneling with a 1D box barrier model (1). The height of the potential barrier (V), the thickness of the potential barrier (L), and the incident, reflected, and transmitted waves are shown. (**C**) The zero-point vibrational energy ($E_0$) of the reactants is lower for O-D than for O-H. As a result, the activation energy is greater for O-D cleavage than for O-H cleavage, thereby causing the kinetic isotope effect (2). (**D**) Chemical reactions of SIRT3-mediated deacetylation of an acetylated lysine residue in the presence of $H_2O$ or $D_2O$.

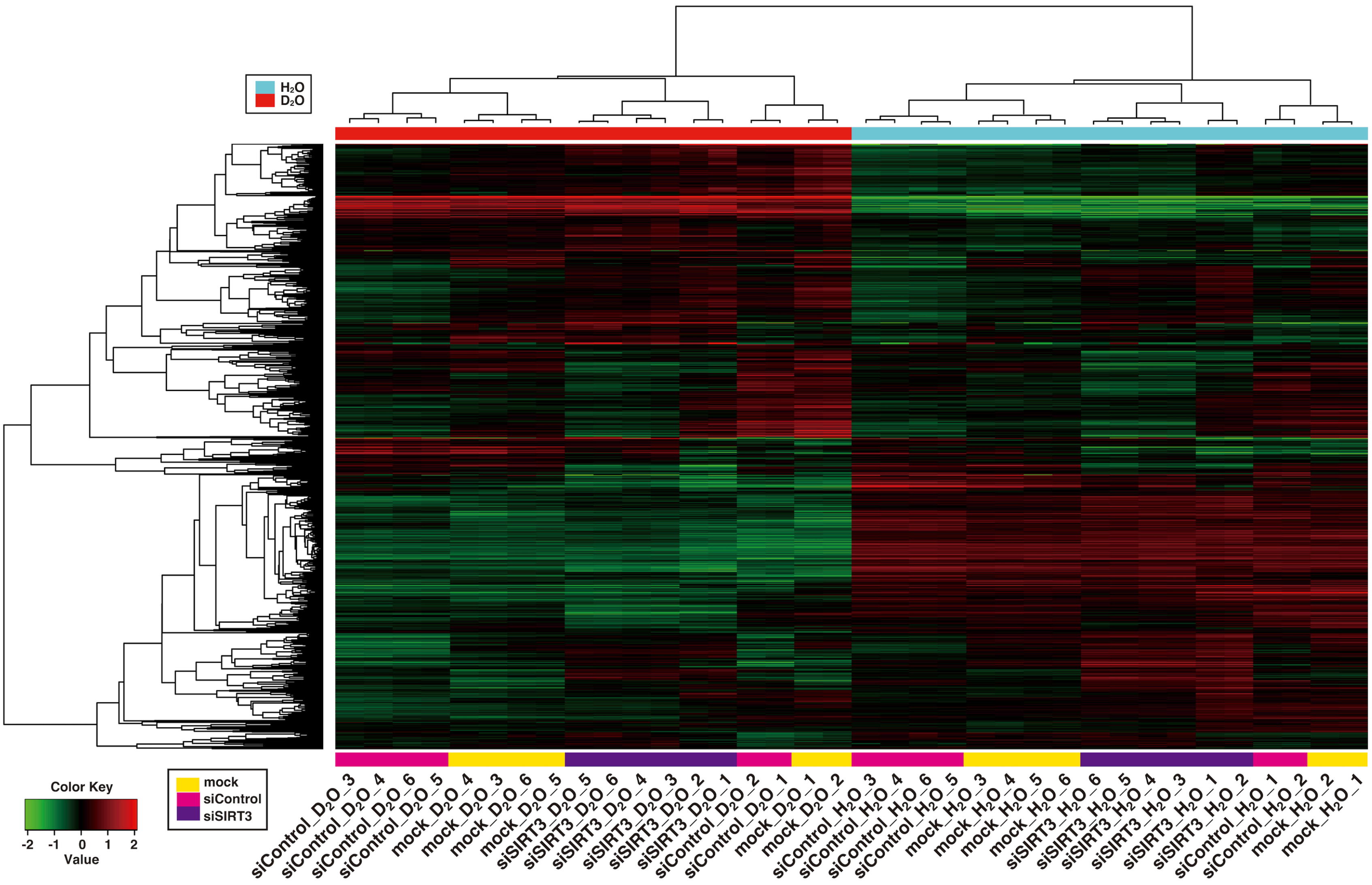
$H_2O$
$D_2O$
Color Key
-2 -1 0 1 2
Value
mock
siControl
siSIRT3
siControl_$D_2O$_3
siControl_$D_2O$_4
siControl_$D_2O$_6
siControl_$D_2O$_5
mock_$D_2O$_4
mock_$D_2O$_3
mock_$D_2O$_6
mock_$D_2O$_5
siSIRT3_$D_2O$_5
siSIRT3_$D_2O$_6
siSIRT3_$D_2O$_4
siSIRT3_$D_2O$_3
siSIRT3_$D_2O$_2
siSIRT3_$D_2O$_1
siControl_$D_2O$_2
siControl_$D_2O$_1
mock_$D_2O$_1
mock_$D_2O$_2
siControl_$H_2O$_3
siControl_$H_2O$_4
siControl_$H_2O$_6
siControl_$H_2O$_5
mock_$H_2O$_3
mock_$H_2O$_4
mock_$H_2O$_5
mock_$H_2O$_6
siSIRT3_$H_2O$_6
siSIRT3_$H_2O$_5
siSIRT3_$H_2O$_4
siSIRT3_$H_2O$_3
siSIRT3_$H_2O$_1
siSIRT3_$H_2O$_2
siControl_$H_2O$_1
siControl_$H_2O$_2
mock_$H_2O$_2
mock_$H_2O$_1

**Fig. S2.**

**Heatmap analysis with hierarchical clustering of the most variable 1,000 genes.** Hela pDR-GFP cells untreated (mock) or transfected with siRNA (siControl or siSIRT3) were cultured in medium made with $H_2O$ or $D_2O$ for 5h, and their RNA samples were subjected to an RNA-seq analysis. Six samples were used for each experimental condition. The heatmap was generated with iDEP96, as described in the Materials and Methods.

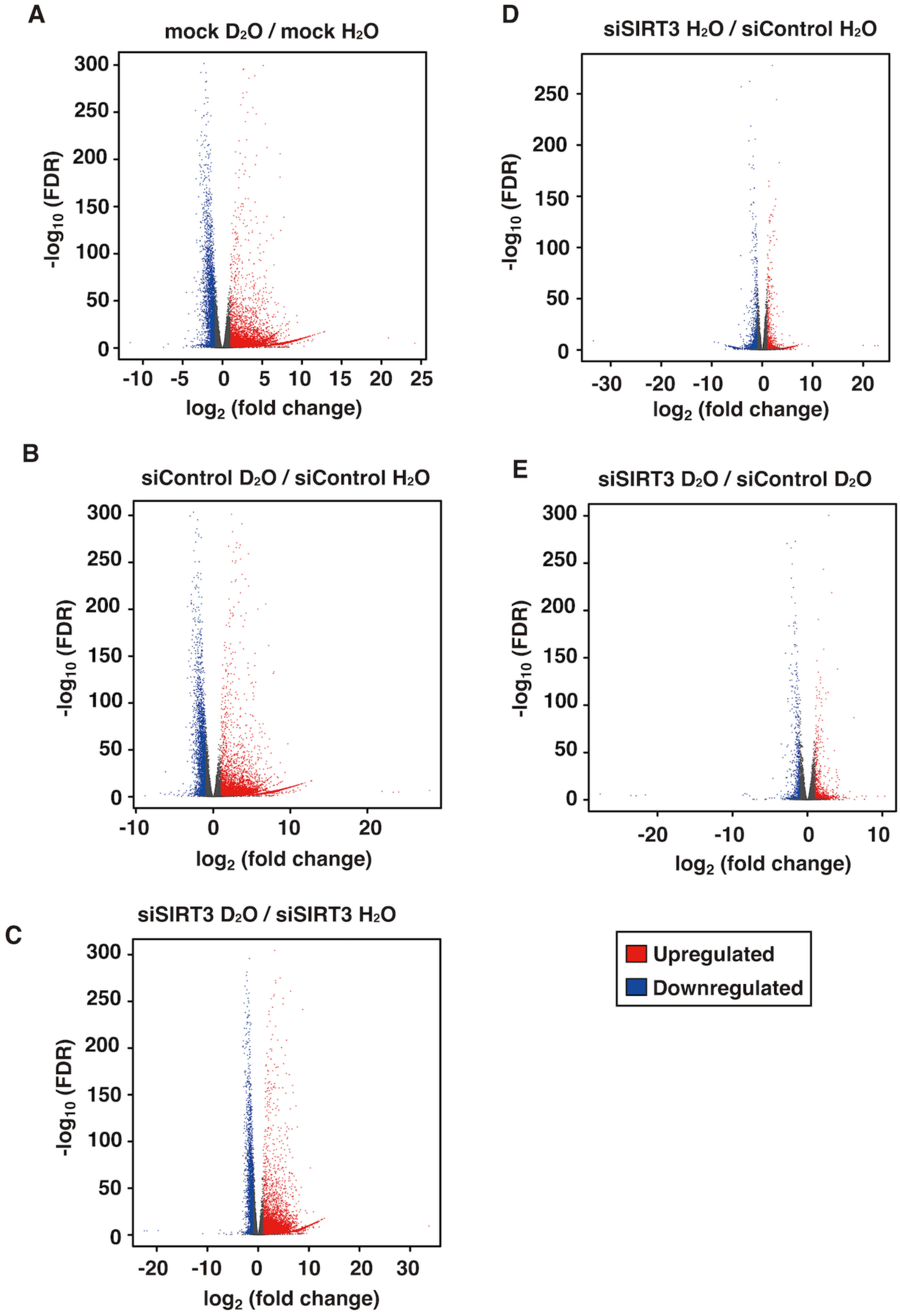
A
mock D₂O / mock H₂O
-log10 (FDR)
log2 (fold change)
B
siControl D₂O / siControl H₂O
-log10 (FDR)
log2 (fold change)
C
siSIRT3 D₂O / siSIRT3 H₂O
-log10 (FDR)
log2 (fold change)
D
siSIRT3 H₂O / siControl H₂O
-log10 (FDR)
log2 (fold change)
E
siSIRT3 D₂O / siControl D₂O
-log10 (FDR)
log2 (fold change)
Upregulated
Downregulated

**Fig. S3.**

**Volcano plots illustrating isotope effects by $D_2O$ on gene expression levels.** The RNA-seq data shown in Fig. S2 were used for a volcano plot analysis with the DEG2 function of iDEP96. The $\log_2$ (fold change) versus -log10 (false discovery rate (FDR)) is shown in each plot.

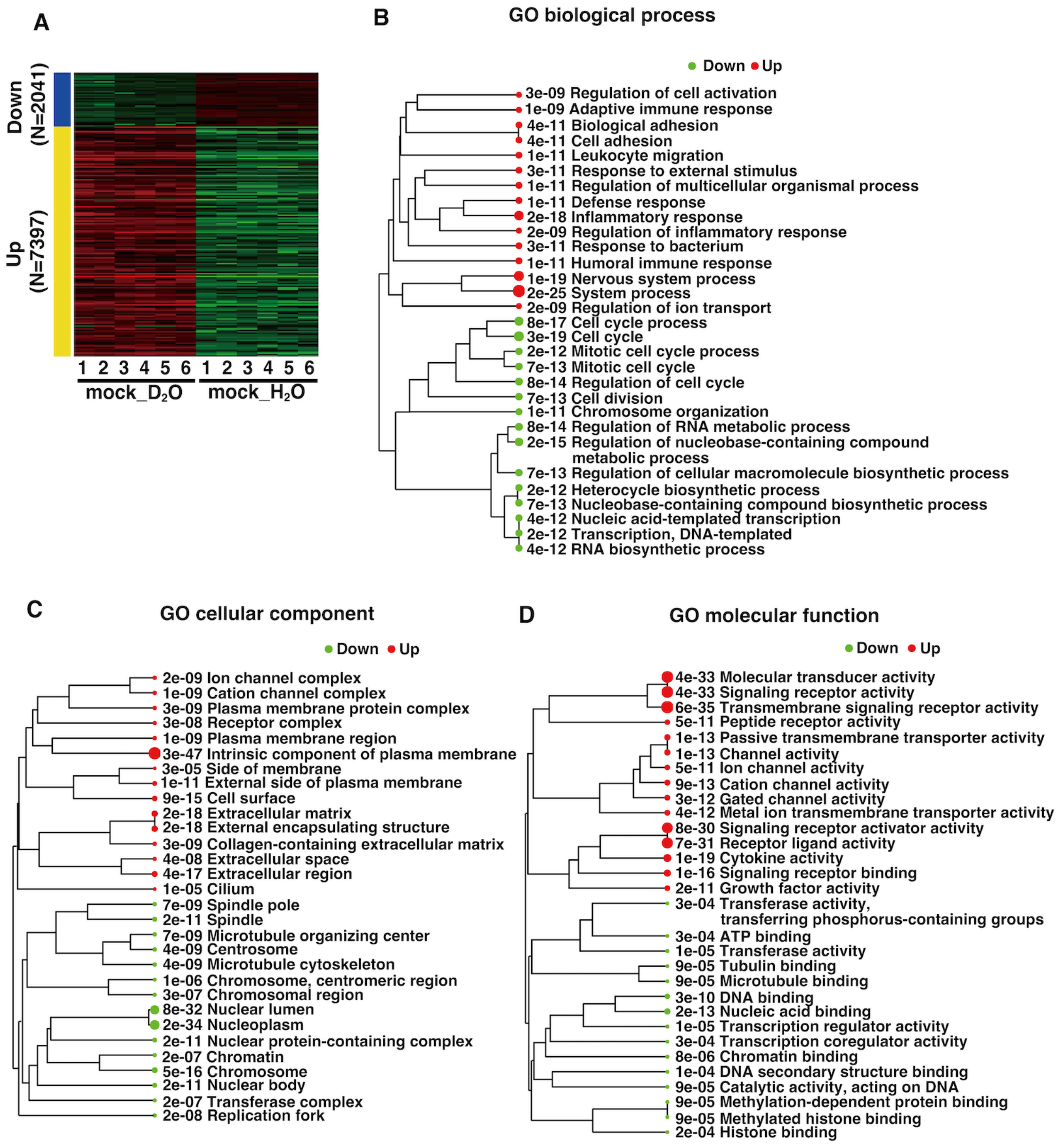


**Fig. S4.**

**Enrichment analysis of gene expression in Hela pDR-GFP cells untreated with siRNA and cultured in the presence of $H_2O$ or $D_2O$.** (**A** to **D**) The RNA-seq data shown in Fig. S2 were used. The RNA-seq data were analyzed with the DEG2 function of iDEP96. Upregulated and downregulated genes are colored red and green, respectively. (**A**) Heatmap analysis of gene expression differences. (**B** to **D**) Enrichment trees. Enrichment pathway analyses were performed for three categories: GO biological process (**B**), GO cellular component (**C**), and GO molecular function (**D**). The false discovery rate (FDR) is shown and is also represented by the size of the circle.

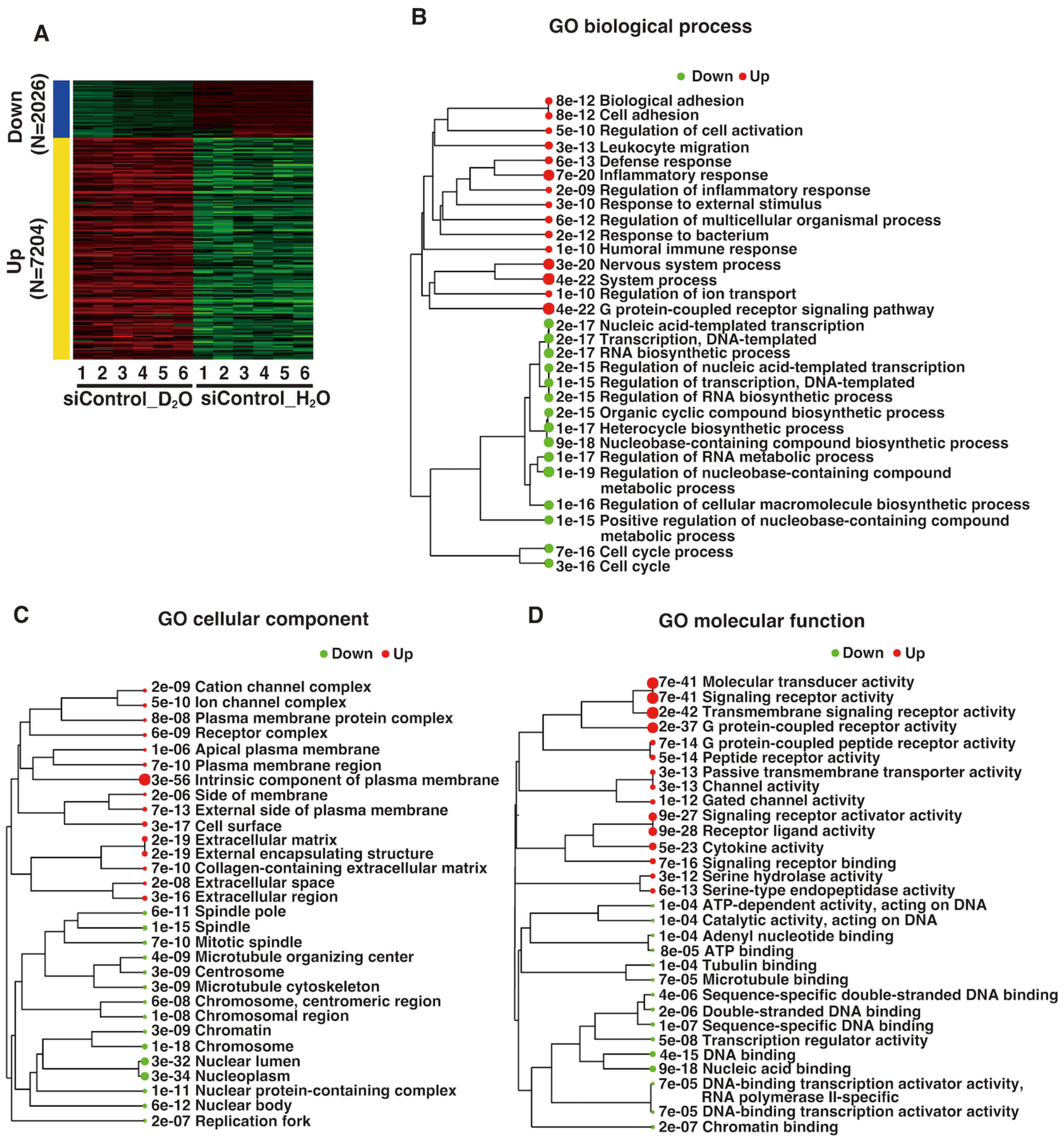


**Fig. S5.**

**Enrichment analysis of gene expression in Hela pDR-GFP cells treated with control siRNA (siControl) and cultured in the presence of $H_2O$ or $D_2O$.** (**A** to **D**) The RNA-seq data shown in Fig. S2 were used. The RNA-seq data were analyzed with the DEG2 function of iDEP96. Upregulated and downregulated genes are colored red and green, respectively. (**A**) Heatmap analysis of gene expression differences. (**B** to **D**) Enrichment trees. Enrichment pathway analyses were performed for three categories: GO biological process (**B**), GO cellular component (**C**), and GO molecular function (**D**). The false discovery rate (FDR) is shown and is also represented by the size of the circle.

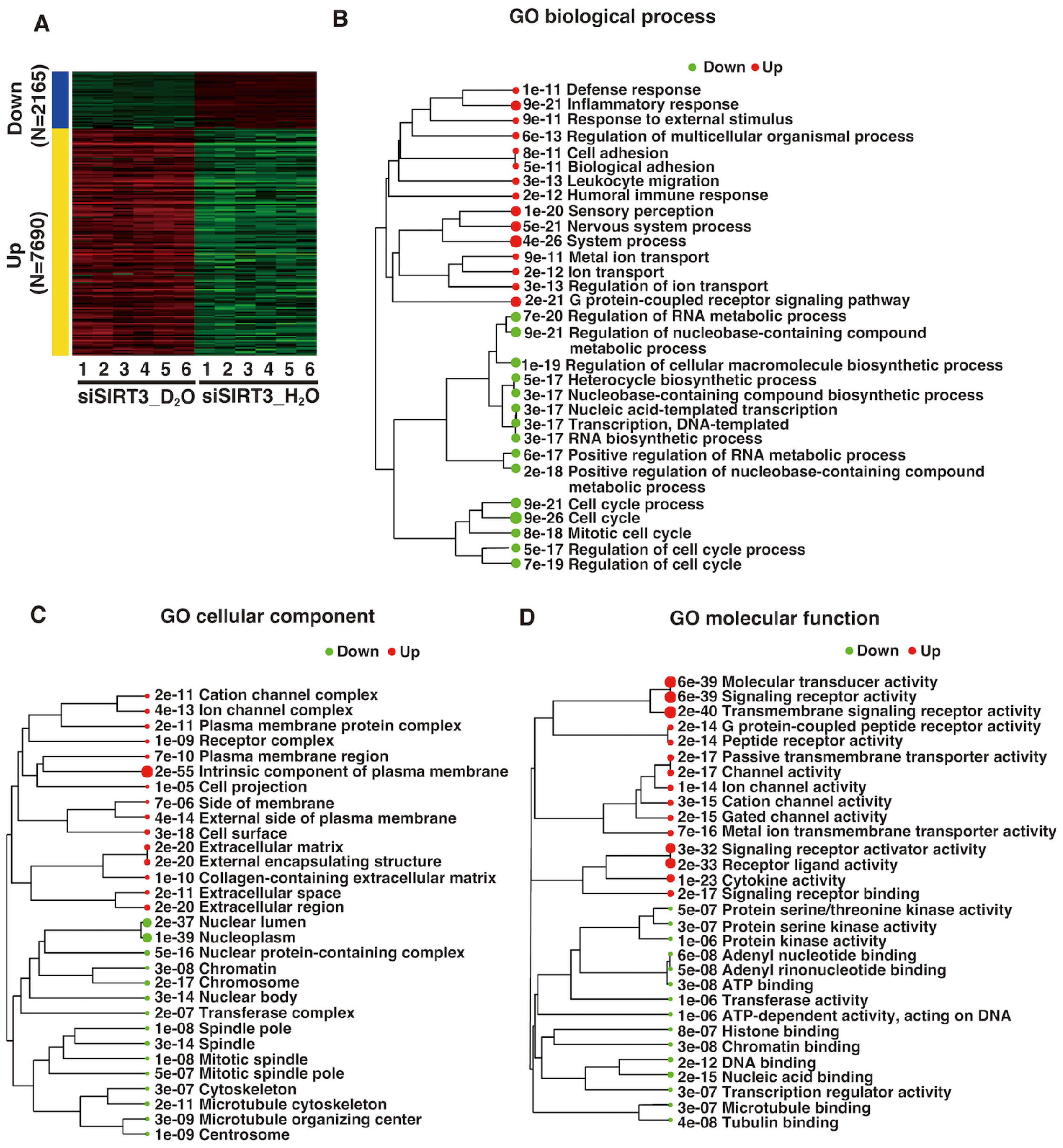


**Fig. S6.**

**Enrichment analysis of gene expression in Hela pDR-GFP cells treated with siRNA against SIRT3 (siSIRT3) and cultured in the presence of $H_2O$ or $D_2O$.** (**A** to **D**) The RNA-seq data shown in Fig. S2 were used. The RNA-seq data were analyzed with the DEG2 function of iDEP96. Upregulated and downregulated genes are colored red and green, respectively. (**A**) Heatmap analysis of gene expression differences. (**B** to **D**) Enrichment trees. Enrichment pathway analyses were performed for three categories: GO biological process (**B**), GO cellular component (**C**), and GO molecular function (**D**). The false discovery rate (FDR) is shown and is also represented by the size of the circle.

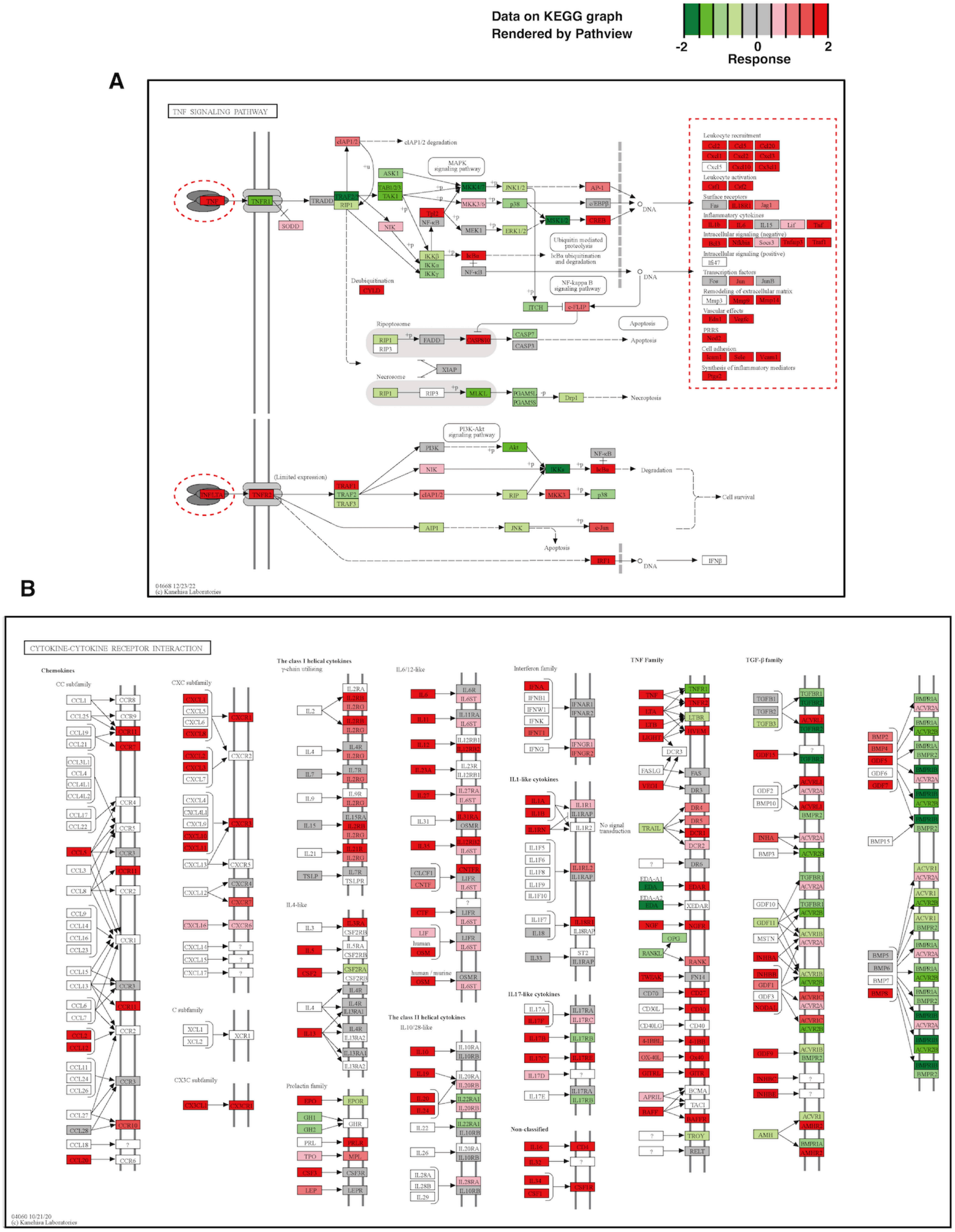


**Fig. S7.**

**Effects of $D_2O$ on expression levels of cellular genes involved in "TNF signaling pathway" and "cytokine-cytokine receptor interaction".** (**A** and **B**) The RNA-seq data shown in Fig. S2 were used. Expression levels of each gene were visualized on KEGG pathway maps of "TNF signaling pathway" (**A**) and "cytokine-cytokine receptor interaction" (**B**), as described in the

Materials and Methods. The red and green colors, according to shading, show an increase and decrease in gene expression, respectively, with $D_2O$ treatment compared to $H_2O$ treatment. (**A**) Characteristic groups of genes with increased expression, such as TNF and cytokines, are surrounded by red dashed lines.

A

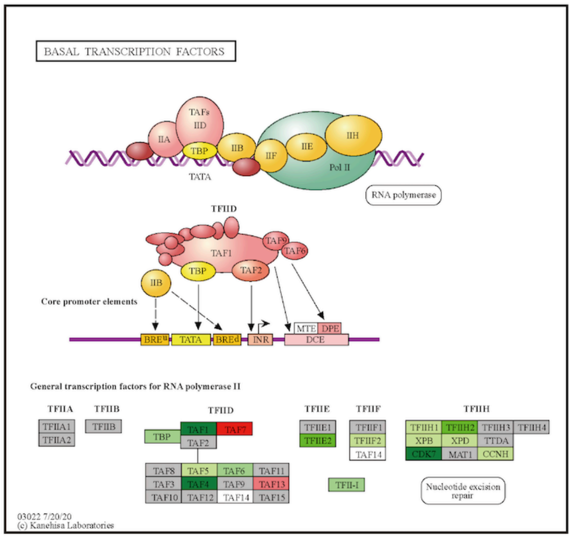
BASAL TRANSCRIPTION FACTORS
RNA polymerase
TFIID
Core promoter elements
General transcription factors for RNA polymerase II
TFIIA
TFIIB
TFIID
TFIIE
TFIIF
TFIIH
Nucleotide excision repair
03022 7/20/20
(c) Kanehisa Laboratories

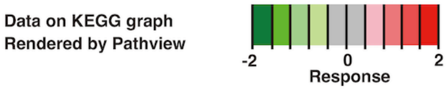
Data on KEGG graph
Rendered by Pathview
-2
0
2
Response

B

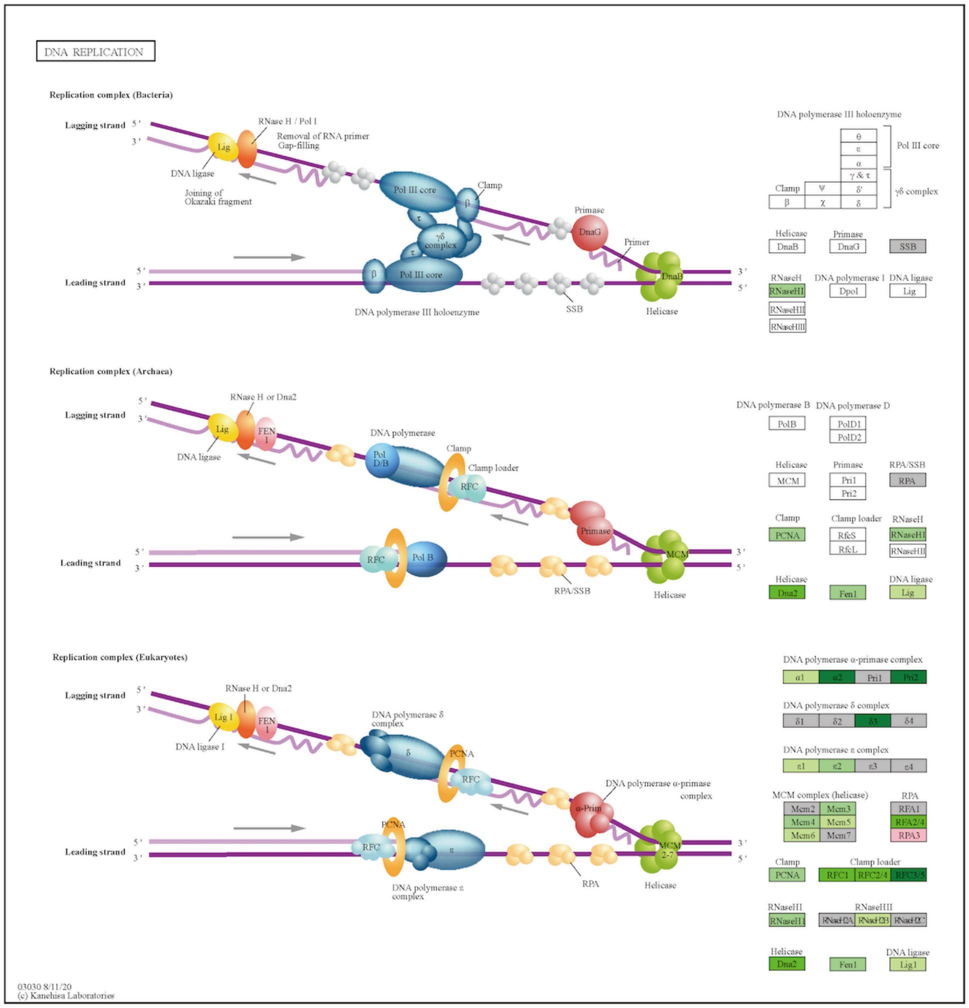
DNA REPLICATION
Replication complex (Bacteria)
Lagging strand
Leading strand
DNA polymerase III holoenzyme
Replication complex (Archaea)
Replication complex (Eukaryotes)
03030 8/11/20
(c) Kanehisa Laboratories

**Fig. S8.**

**Effects of $D_2O$ on expression levels of cellular genes involved in "basal transcription factors" and "DNA replication".** (**A** and **B**) The RNA-seq data shown in Fig. S2 were used. Expression levels of each gene were visualized on KEGG pathway maps of "basal transcription factors" (**A**) and "DNA replication" (**B**), as described in the Materials and Methods. The red and green colors, according to shading, show an increase and decrease in gene expression, respectively, with $D_2O$ treatment compared to $H_2O$ treatment.

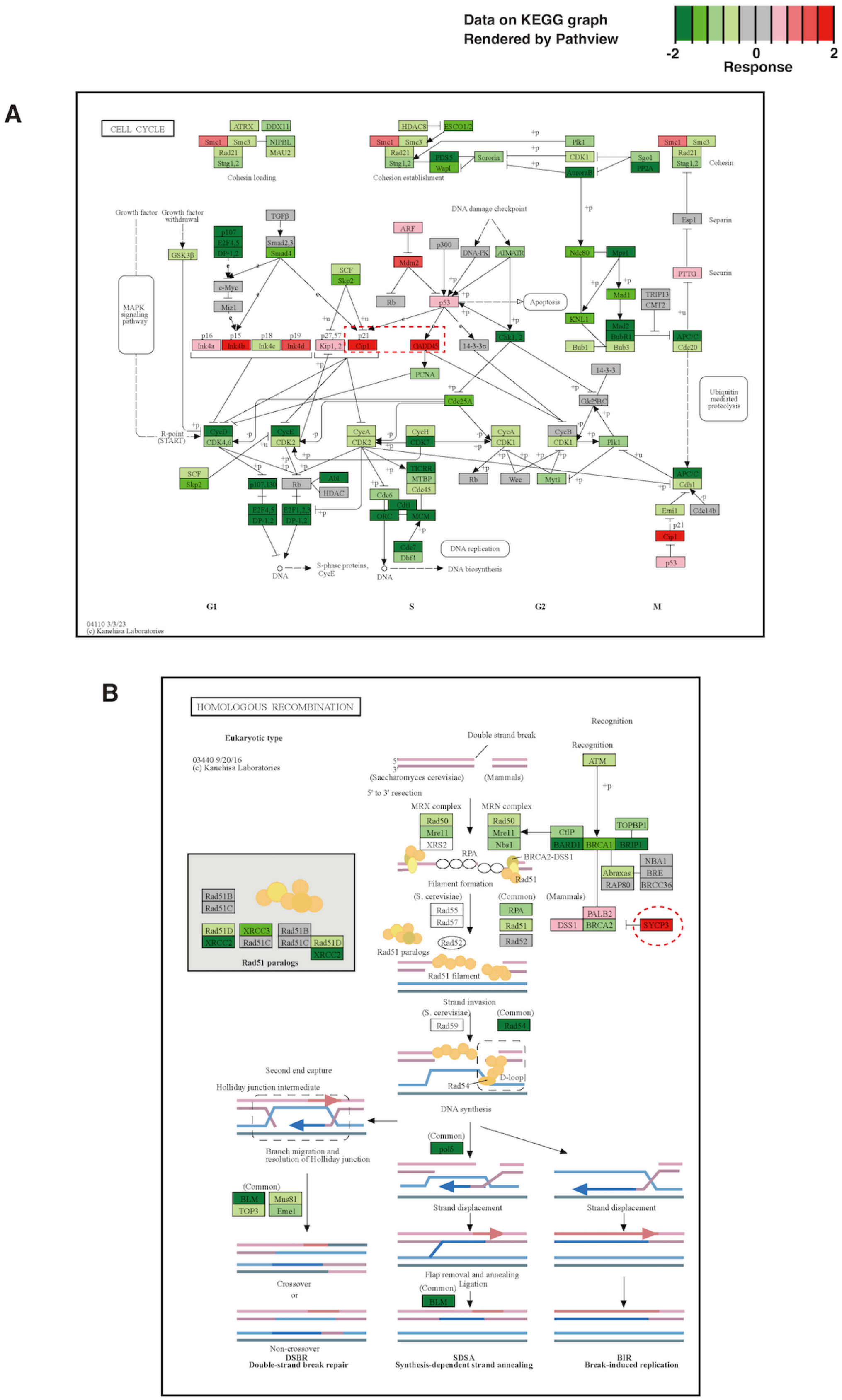
Data on KEGG graph
Rendered by Pathview
-2
0
2
Response
A
CELL CYCLE
Cohesin loading
Cohesion establishment
Cohesin
DNA damage checkpoint
Separin
Securin
Apoptosis
MAPK signaling pathway
Growth factor
Growth factor withdrawal
R-point (START)
Ubiquitin mediated proteolysis
DNA replication
S-phase proteins, CycE
DNA biosynthesis
DNA
G1
S
G2
M
04110 3/3/23
(c) Kanehisa Laboratories
B
HOMOLOGOUS RECOMBINATION
Eukaryotic type
03440 9/20/16
(c) Kanehisa Laboratories
Recognition
Double strand break
(Saccharomyces cerevisiae)
(Mammals)
5' to 3' resection
MRX complex
MRN complex
Filament formation
Rad51 paralogs
Rad51 filament
Strand invasion
D-loop
DNA synthesis
Second end capture
Holliday junction intermediate
Branch migration and resolution of Holliday junction
Strand displacement
Flap removal and annealing
Ligation
Crossover
or
Non-crossover
DSBR
Double-strand break repair
SDSA
Synthesis-dependent strand annealing
BIR
Break-induced replication

**Fig. S9.**

**Effects of $D_2O$ on expression levels of cellular genes involved in "cell cycle" and "homologous recombination".** (**A** and **B**) The RNA-seq data shown in Fig. S2 were used. Expression levels of each gene were visualized on KEGG pathway maps of "cell cycle" (**A**) and "homologous recombination" (**B**), as described in the Materials and Methods. The red and green colors, according to shading, show an increase and decrease in gene expression, respectively, with $D_2O$ treatment compared to $H_2O$ treatment. Characteristic groups of genes with increased expression, such as inhibitors of cell cycle or HR repair, are surrounded by red dashed lines.

**A**

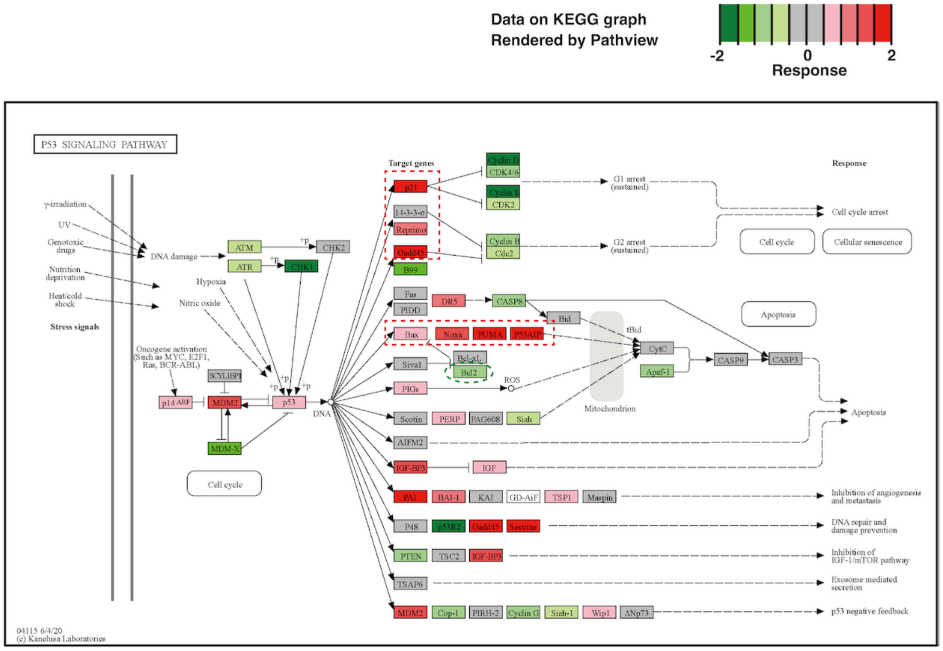


**B**

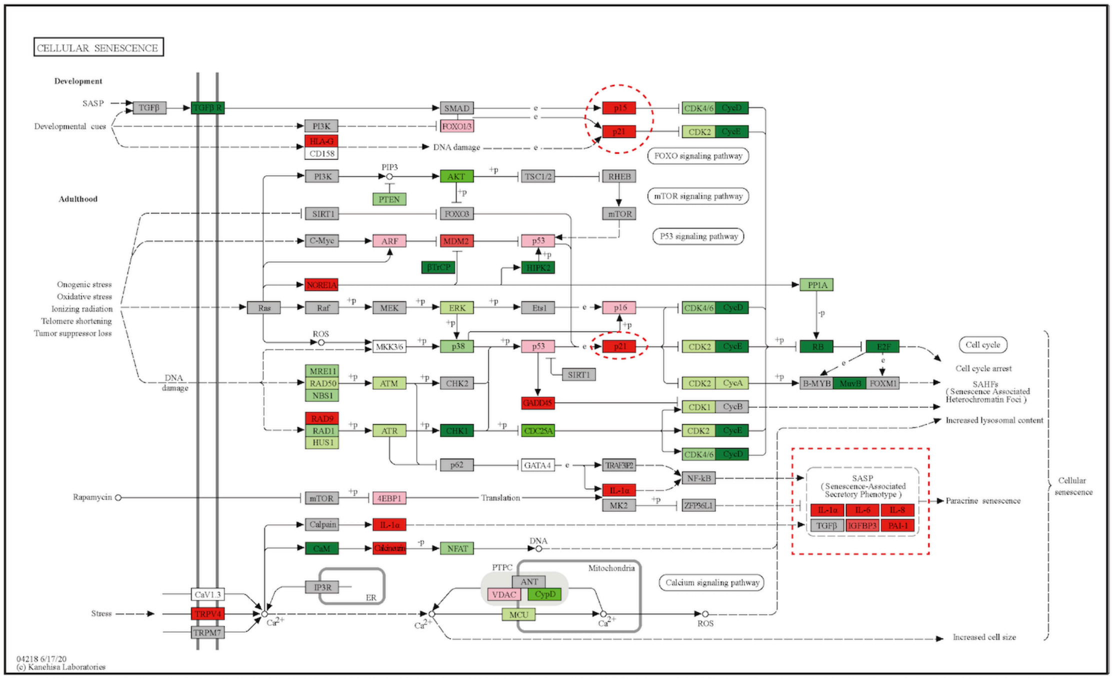


**Fig. S10.**

**Effects of $D_2O$ on expression levels of cellular genes involved in "p53 signaling pathway" and "cellular senescence".** (**A** and **B**) The RNA-seq data shown in Fig. S2 were used. Expression levels of each gene were visualized on KEGG pathway maps of "p53 signaling pathway" (**A**) and "cellular senescence" (**B**), as described in the Materials and Methods. The red and green colors, according to shading, show an increase and decrease in gene expression, respectively, with $D_2O$ treatment compared to $H_2O$ treatment. Characteristic groups of genes with increased or decreased expression are surrounded by red or green dashed lines, respectively.

**Data S1. (separate file)**

**The obtained read counts data from RNA-seq experiments.** The CSV file of the data S1 used for the iDEP96 analysis. Figs. S2-S10 were obtained with data S1.

**Data S2. (separate file)**

**The experimental design file for the iDEP96 analysis.** The CSV file of data S2 used for the iDEP96 analysis with the CSV file of data S1.
